# A refined method to monitor arousal from hibernation in the European hamster

**DOI:** 10.1101/2020.04.20.009712

**Authors:** Fredrik A. S. Markussen, Vebjørn J. Melum, Béatrice Bothorel, David G. Hazlerigg, Valérie Simonneaux, Shona H. Wood

## Abstract

**Background:** Hibernation is a physiological and behavioural adaptation that permits survival during periods of reduced food availability and extreme environmental temperatures. This is achieved through cycles of metabolic depression and reduced body temperature (torpor) and rewarming (arousal). Rewarming from torpor is achieved through the activation of brown adipose tissue (BAT) associated with a rapid increase in ventilation frequency. Here, we studied the rate of rewarming in the European hamster (*Cricetus cricetus*) by measuring both BAT temperature, core body temperature and ventilation frequency.

**Results:** Temperature was monitored in parallel in the BAT (IPTT tags) and peritoneal cavity (iButtons) during hibernation torpor-arousal cycling. We found that increases in brown fat temperature preceded core body temperature rises by about 48 min, with a maximum re-warming rate of 20.9°C*h^−1^. Re-warming was accompanied by a significant increase in ventilation frequency. The rate of rewarming was slowed by the presence of a spontaneous thoracic mass in one of our animals. Core body temperature re-warming was reduced by 6.2°C*h^−1^ and BAT rewarming by 12°C*h^−1^. Ventilation frequency was increased by 77% during re-warming in the affected animal compared to a healthy animal. Inspection of the position and size of the mass indicated it was obstructing the lungs and heart.

**Conclusions:** We have used a minimally invasive method to monitor BAT temperature during arousal from hibernation illustrating BAT re-warming significantly precedes core body temperature re-warming, informing future study design on arousal from hibernation. We also showed compromised re-warming from hibernation in an animal with a mass obstructing the lungs and heart, likely leading to inefficient ventilation and circulation.

## Background

Hibernation is a physiological and behavioural adaptation that permits survival during periods of reduced food availability and extreme environmental temperatures. This is achieved through cycles of metabolic depression characterised by reduced body temperature (torpor) and rewarming (arousal). Entrance into torpor precisely is controlled by decreases in heart rate, ventilation frequency and oxygen consumption [1, 2]. Arousal occurs in a coordinated manner with increased ventilation frequency and oxygen consumption subsequently followed by heart rate, blood pressure and core body temperature rise [1]. Rewarming from torpor is achieved through the activation of subcutaneous brown fat reserves on top of the scapulae (classical BAT) and within the intra-scapular region (intra-scapular BAT) [3]. The primary function of brown fat is heat generation through non shivering thermogenesis [4], which has a high energy (oxygen) demand [5, 6]. During arousal from torpor temperature increases in brown fat correlate with increased oxygen consumption (respiration rate) in the brown bat (*Eptesicus fuscus*) and the golden-mantled ground squirrel (*Callospermophilus lateralis*), and precede rectal or muscle temperature increases [7–10].

Activation of brown fat is thought to originate in the thermo-sensing regions in the hypothalamus coupled to a sympathetic nervous pathway, which activates beta adrenergic receptors and in turn the brown fat mitochondria [11]. The thermogenic capacity of brown fat comes from the use of proton leak (uncoupling) in mitochrondria as opposed to the coupled oxidative phosphorylation pathway which produces ATP [12]. The uncoupled pathway depends on the BAT-specific expression of the uncoupling protein, UCP1 transporter in the mitochondrial membrane. Initiation of this process requires good oxygen/energy supply to the BAT, and a marked increase in ventilation is an early event in the rewarming process [13]. The heat generated from BAT rewarms the anterior of the animal first, and then increases in heart rate and circulation are required to warm the rest of the animal and raise core body temperature (Tb). Subsequently, shivering thermogenesis is initiated to help the animal to reach normal body temperature (euthermy) quickly [5, 6, 14]. In Syrian hamsters (*Mesocricetus auratus*) the initiation of shivering thermogenesis can only occur once the body temperature is above 16□ [5] and in the marmot (*Marmota marmota*) shivering is only observed when the subcutaneous BAT temperature reaches 16□ [4]. These data highlight the importance of non-shivering thermogenesis by BAT in the initial stages of the re-warming process.

Tb monitoring by iButtons implanted into the intraperitoneal cavity is the standard method used to monitor torpor arousal cycling during hibernation [15–18], and while it is known that BAT activation is the first stage in the re-warming process it is rarely monitored and good characterisation of the relationship between BAT and core body temperature during hibernation is lacking. Here, we use a minimally invasive method to monitor both BAT temperature (T_BAT_) and Tb in a well-established hibernation model, the European hamster (*Cricetus cricetus*) [19–21]. We demonstrate that increases in T_BAT_ precede increases in Tb and that the ventilation frequency correlates with the rate of T_BAT_ re-warming. Furthermore, one animal with a thoracic mass showed impaired ventilation which led to a marked slowing of the re-warming process.

## Results

### Re-warming in brown fat compared to the core body

We placed Implantable Programmable Temperature Transponder (IPTT) tags subcutaneously to measure the T_BAT_. In the same animals we surgically implanted iButtons into the intraperitoneal cavity to monitor Tb. To initiate the preparation for hibernation we transferred animals from long photoperiod (14L:10D, 14 hours of light per 24 hours) and 22□ (LP22), to short photoperiod (10L:14D, 10 hours of light per 24 hours) and 22□ (SP22). After 8 weeks, we reduced the room temperature to 10□ (SP10, Figure 1A). All animals showed a hibernation phenotype within 4 weeks of transfer to SP10, which was preceded by “test drops” in Tb of approximately 10 to 15□ below euthermy before initiating the multi-day torpor-arousal cycling characteristic of the hibernation season (Figure 1B). Once hibernating, the European hamster drops its Tb to near ambient room temperature for multiple days (approximately 25□ below euthermy) and arouses at intervals returning to euthermy (Figure 1B).

**Figure 1:**
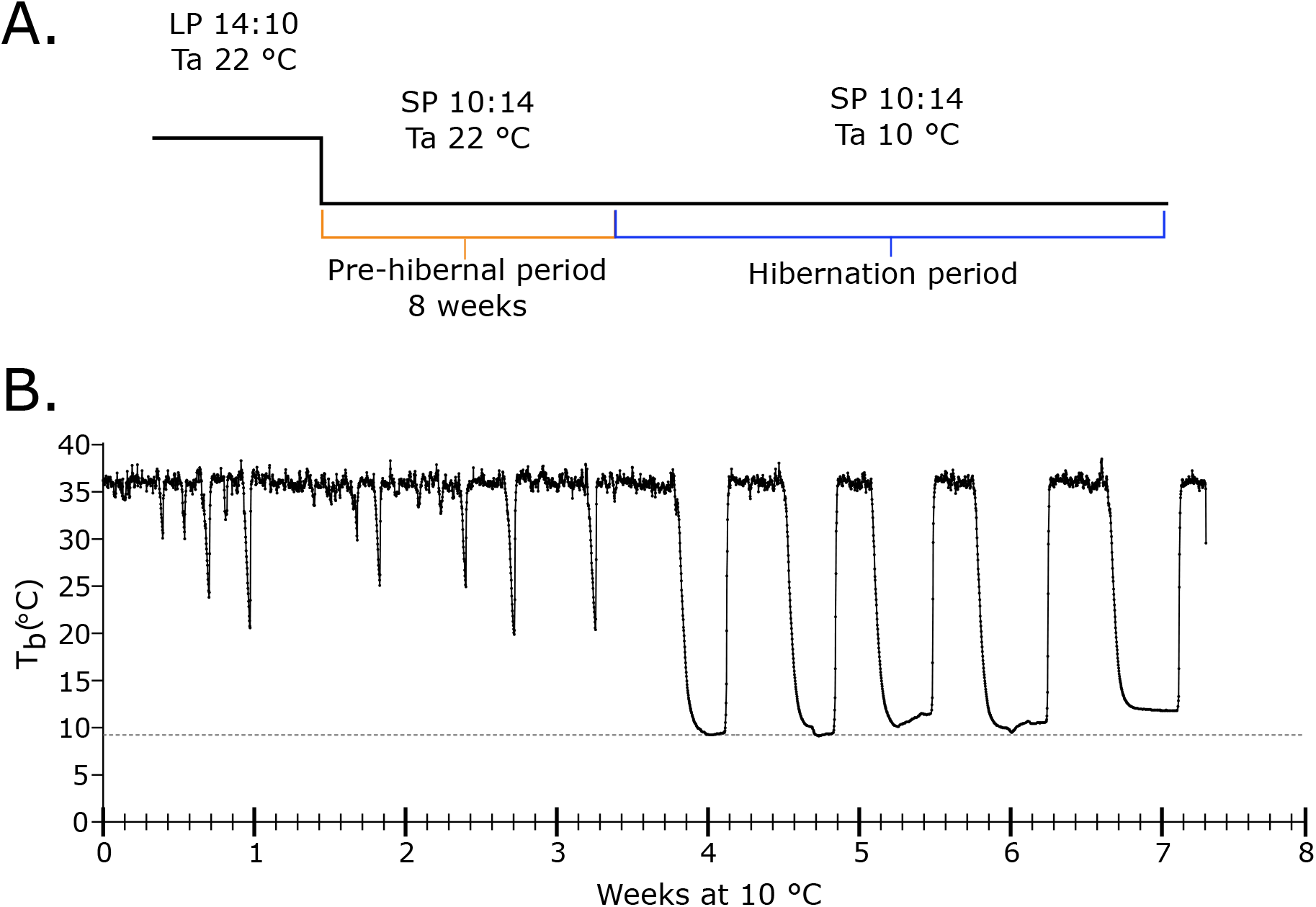
European hamster displays hibernation after exposure to short photoperiod and low ambient temperature. A.) Study design to induce the hibernation phenotype. Animals were transferred from long photoperiod (14 hours of light per day, 14:10) and 22°C to short photoperiod (10 hours of light per day, 10:14) for 8 weeks. The ambient temperature (Ta) was lowered to 10°C, a temperature of 9.6 ± 1°C was observed. B.) Core body temperature (Tb) measurement from a European hamster displaying torpor-arousal cycling. The grey dotted line indicates the average ambient temperature (Ta) over the experiment.

We confirmed the initiation of the hibernation season and torpor-arousal cycling for at least 2 weeks before we started monitoring T_BAT_ during the periodic arousals. These data were time matched with Tb from the iButtons. Arousal events were recorded from five individuals (3 females, 2 males). We observed that both T_BAT_ and Tb show similar sigmoidal curve trajectories until euthermy is reached (Figure 2A, non-linear asymmetric sigmoidal model, T_BAT_: r^2^= 0.9778; Tb: r^2^= 0.9878). We show that BAT re-warming precedes subsequent core body re-warming by 48.6 minutes (CI: 45.4-51.7 minutes) (Figure 2A). The mean ventilation frequency (VF) for the 5 individuals was calculated for each quartile of re-warming showing a clear linear increase in VF during re-warming which correlated with T_BAT_ (Figure 2A, one-way ANOVA p<0.0001, R^2^=0.86). Using visual assessment, the onset of shivering in muscles adjacent to the BAT during arousal was noted, an average BAT temperature of 15□ (CI: 14.2-16.8□) was recorded at the onset of shivering (Figure 2B).

**Figure 2:**
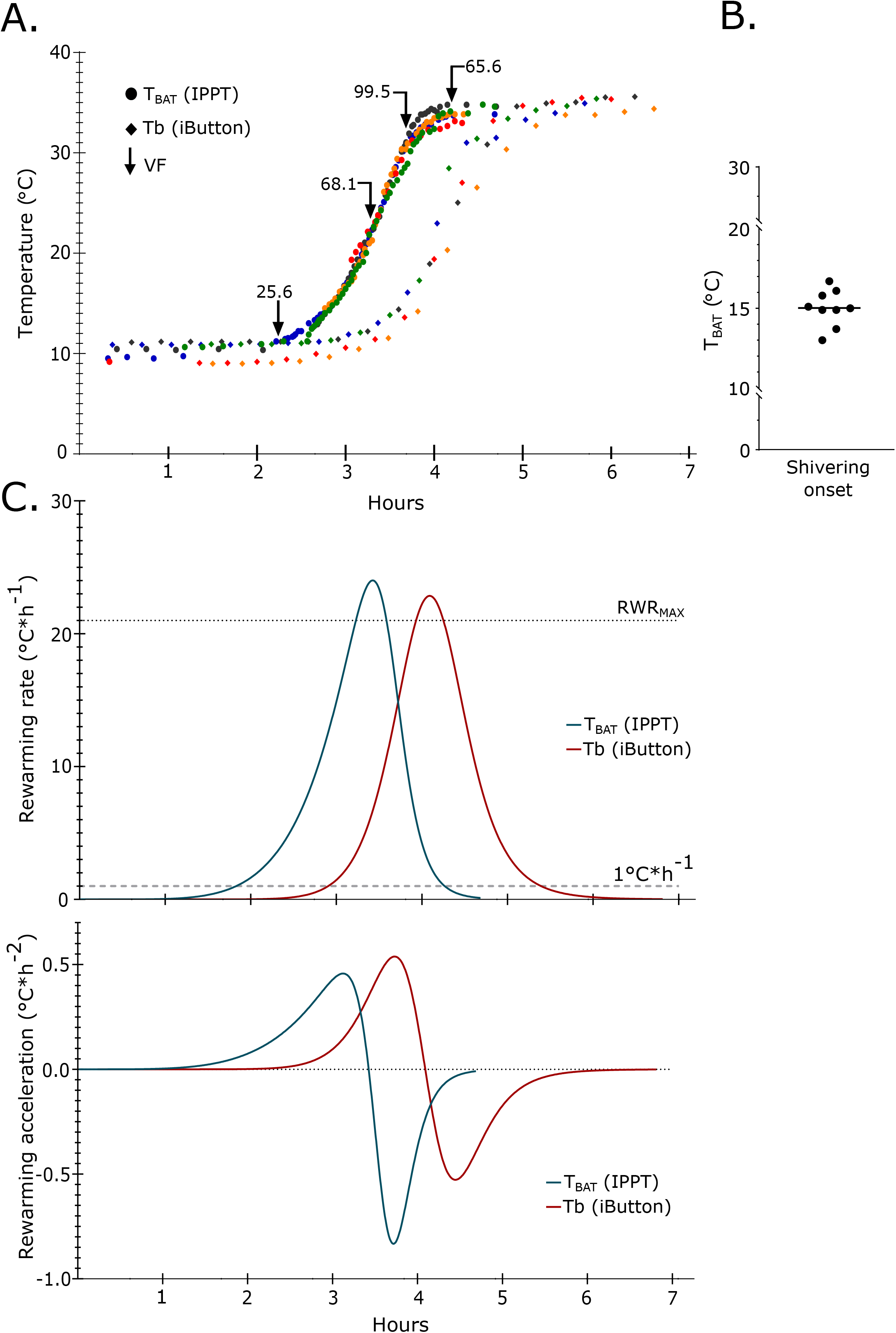
Brown adipose tissue re-warming precedes core body temperature increases. A) Core body temperature (Tb – iButton, diamond symbol) and BAT temperature recordings (T_BAT_ - IPTT tag, circle symbol) from 5 individuals during an arousal from torpor (each individual represented by a different colour). The mean ventilation frequency (VF) at each quartile of arousal, as defined by a non-linear curve fit for all individuals, is indicated by black arrows. B) BAT temperature recording corresponding to onset of BAT adjacent skeletal muscle shivering. Observations and recordings were done in 9 separate arousal events in 5 different animals. C.) Five arousal events synchronised in time at 23.5°C showing the modelled 1st and 2nd derivatives. First derivate graph describes rewarming rate (RWR) and second derivate graph the rewarming acceleration. Mean maximum rewarming rate (RWRmax, best fit linear regression) is similar for both Tb and TBAT at 21.0 C∗h −1 (CI: 15.4- 26.5°C∗h −1) and 20.9°C∗h −1 (CI: 20.3-21.4°C∗h −1), respectively.

We calculated the maximum re-warming rate (RWR_MAX_) by analysing the linear phase of the sigmoidal curve and fitting a best fit regression line (Figure 2C). We found that RWR_MAX_ was similar for BAT (RWR_MAX_ = 20.9□*h^−1^; 95% confidence interval (CI): 20.3□ to 21.5□) and core body (RWR_MAX_ = 21.0□*h^−1^; CI: 15.4□ – 26.5□) (Figure 2C), however Tb RWR_MAX_ showed more individual variance. We then calculated the rate of change in re-warming (acceleration), or how fast RWR_MAX_ is achieved, by using the derivatives of the sigmoidal curve in the exponential and asymptotic phases (Figure 2C). We found the initiation of re-warming (exponential phase) is delayed by 66 minutes in the core body compared to the BAT. However, the time to reach RWR_MAX_ is less in the core body (59 minutes) compared to the BAT (83 minutes)(Figure 3C).

**Figure 3:**
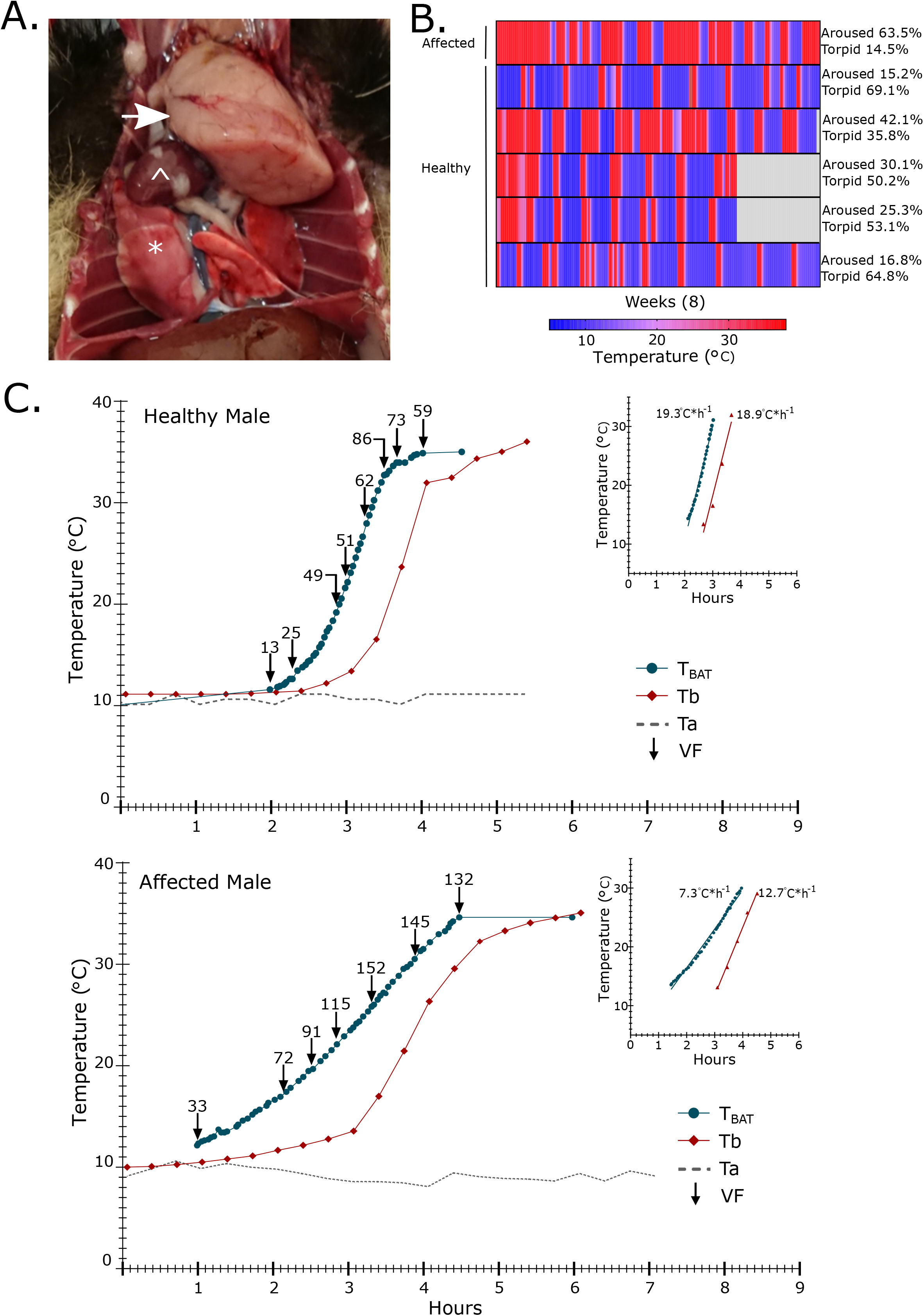
Rewarming from torpor is compromised in an animal with a thoracic mass. A.) Post-mortem image of exposed thoracic cavity of a European hamster with a thoracic mass (indicated by the white arrow). For reference, white * is placed on lungs and ^ on heart. B.) Heat-map of core body temperature recorded by iButtons over the course of the experiment. Each line is an individual animal, the first animal has the thoracic mass. The percentage time spent above 35.5°C (Euthermic – aroused) and below 12°C (Torpid) is indicated, the remainder is time spent between 35 to 12 °C (entering or arousing from torpor) for each individual. Grey indicates no data was recorded in this period. C.) Core body temperature (Tb), BAT temperature (T_BAT_) and ventilation frequency (VF) changes during arousal from torpor in a healthy male and an affected male with a thoracic mass. Grey dotted line shows room temperature (Ta). Inset graph shows the linear portion of the arousal that was used to calculate maximum re-warming rate (RWR_max_) in °C per hour.

### Brown fat re-warming is compromised by the presence of a spontaneous thoracic mass

We identified one individual in our study with a spontaneous thoracic mass (Figure 3A). When comparing the amount of time spent torpid or aroused the affected animal spent 63.5% aroused and 14.5% torpid, compared to an average of 25.9% aroused and 54.6% torpid for the healthy animals (Figure 3B). The RWR_max_ of brown fat was reduced by 12□*h^−1^ relative to a representative healthy animal (Figure 3C). The RWR_max_ of the core body was less compromised showing a reduction of 6.2□*h^−1^. Ventilation frequency increases still correlated with T_BAT_ (Healthy: pearson r= 0.899, p-value= 0.002, affected: pearson r= 0.873, p-value= 0.002) but the affected animal showed a 77 % higher maximum ventilation frequency compared to a healthy animal (Figure 3C). BAT re-warming precedes core body re-warming by 53.9 minutes in the affected animal, this is outside the confidence intervals for the healthy animals, suggesting that the efficiency of BAT re-warming of the core is compromised.

## Discussion

Our study has used a minimally invasive method to measure BAT temperature during hibernation. We show that there is a significant correlation between T_BAT_ and Tb, with T_BAT_ increases preceding that of Tb. Compared to intraperitoneal implantation of iButtons the IPTT tags offer a minimally invasive method to monitor temperature, furthermore there is a significant advantage to monitoring BAT temperature in hibernation studies instead of core body temperature because the earliest arousal events can be detected almost immediately.

The increases in T_BAT_ and Tb are both correlated with increased ventilation frequency. Hyperventilation has been previously observed in the thirteen-lined ground squirrel but BAT temperature was not studied [13]. Our T_b_ RWR_max_ are similar to previous observations in rodents and European hamsters [22] but no data on BAT temperature is available.

Re-warming efficiency of the BAT in the affected animal was only 38% of that in the healthy animal, whereas the rewarming efficiency of the core body was 67% of that in the healthy animal. We also observed a greatly increased ventilation frequency compared to a healthy animal. We suggest that the size and position of the thoracic mass obstructed the lungs, reducing ventilation volume, thereby compromising the animal’s ability to deliver the necessary oxygen to support the activity of BAT. This probably resulted in increased ventilation frequency in an attempt to meet tissue demands. It has been observed that the early arousal events show a reliance on increased ventilation frequency and that increases in heart rate occur later as peripheral circulation increases [13]. The generally lower re-warming efficiency in both BAT and core body might be interpreted as evidence for compromised cardiac function leading to reduced circulation efficiency. In support of a decreased rate of re-warming of the BAT and core body due to decreased circulation we see re-warming rates in the thymus tumour animal comparable to that of the much larger Marmot (5000g compared to 500g) [22].

The discrepancy between BAT re-warming efficiency and core body re-warming is intriguing, and possibly relates to complicated relationship and order of events in arousal which start with increased ventilation to increase oxygen availability, followed by vasodilation, and increased heart rate, to re-perfuse the circulation and re-warm the whole animal. Low oxygen availability increases vasodilation [23] therefore in the affected animal lower oxygen availability may have increased vasodilation leading to a compensated re-warming efficiency of the core body. Vasodilation would also benefit the BAT by supplying more oxygen in the blood. In support of this the affected animal in the initial stages of BAT re-warming from 10 to 15□ appears to show a gentler slope compared to 20 to 30□ (Figure 3C), indicating a lower efficiency in the early arousal stage which would not benefit from vasodilation, this distinction is not seen in the healthy animals. Another explanation for the discrepancy between BAT and core body re-warming efficiency may be the additional contribution of shivering thermogenesis to core body temperature increases.

We propose that the thoracic mass observed in this study is a thymus tumour, and whilst, thymus tumours are rare they are reported in domestic, laboratory and wild contexts [24]. The thymus is a lymphoid organ involved in T-cell maturation, it is located in front of the heart and found in all vertebrates (reviewed in: [25]). However, without histology we cannot ascertain whether the observed thoracic mass is a neoplasia, granuloma or an abscess, nevertheless thymus tumours have been previously histologically characterised in this laboratory colony of European hamsters, the size, position and incidence we observe closely matches previous observations [26]. This previous characterisation also noted a close match human thymic epithelial tumours [26]. Human thymus tumour cases often present with tumours that obstruct the heart and lungs, similar to our observations. Also in humans, an association of thymus tumour with hypothermia/defective re-warming has been reported twice [27, 28]. Impressively in one of the cases the patient experienced a body temperature between 32.8□ to 35□ (Ambient room temperature: 22□) [27]. However, the cause of hypothermia in these cases is unknown.

## Conclusions

In conclusion we have used temperature logging in BAT and intraperitoneal cavity to study progression of arousal in hibernating hamsters. We also showed compromised re-warming from hibernation in an animal with a mass obstructing the lungs and heart, likely leading to inefficient ventilation and circulation.

## Methods

### Animals and ethics statement

European hamsters were bred from stock animals at the Chronobiotron, an animal facility dedicated to the study of biological rhythms. Animals were housed in an environment controlled room in separate cages according to legislations provided in European Commission directive 2010/63/EU. Specifically, animals were housed individually due to aggressive behavior in static type III macrolon cages (Bioscope, Ammendinger, Germany). Environmental enrichment was provided by cardboard nesting material and gnawing sticks.

They were provided with *ad libitum* access to food (Safe^®^005 diet, Safe, Augy, France) and water throughout the study period. The animals were kept under a long photoperiod (14L:10D) and at 22□ (LP22) until the start of the experiment. All animals were 1.5 years old at the start of the experiment. To initiate the preparation for hibernation (pre-hibernal period) the animals were transferred to a short photoperiod (10L:14D) at 22□ (SP22). After 8 weeks, the temperature was lowered to 10□ (SP10), all animals exhibited torpor-arousal cycling, and were kept in this condition to the end of the experiment (10 weeks, Figure 1A). 48 animals (22 male, 26 female) underwent hibernation. Post hoc analysis of temperature data from six hamsters (3 male and 3 female) are included in this study, the remaining hamsters are part of another study. This is a longitudinal study design monitoring individual animals over 8 weeks during torpor arousal cycling. All hamsters were euthanized at the end of the study by a combination of isoflourane, Zoletil/Xlyazine and decapitation. The experimental procedure was validated by a local ethical committee and further validated by the Ministry of Higher Education, Research and Innovation (APAFIS#21424-2019070219421923 v3).

### Calibration of IPTT tags

The accuracy of each IPTT tag (BMDS IPTT-300^®^) was verified from manufacturer to be ±1□ of actual temperature in the range between 21 and 30□. During torpor-arousal the tags would measure between 10-38□ therefore to ensure accuracy between the iButtons and IPTT tags we created an individual calibration curve for each IPTT tag allowing for *post hoc* correction at a 0.1□ resolution. A similar method was used by Wacker et al [29]. Each tag was calibrated using a water bath containing two iButtons (thermochron DS1922L, Maxim integrated) set to 16-bit resolution (0.0625□) and sampling rate of 2 seconds, the IPTT tag was scanned every 2-3 seconds. The water bath was then placed on a heating plate with a magnetic stirrer and heated, the temperature range recorded was 5□ to 39□. The resulting data was plotted in Graphpad Prism v8.0 and a regression analysis was done to allow for calibration of the IPTT tag recordings.

### iButton and IPTT tag surgery

Surgeries were performed on each animal to implant iButtons and IPTT tags. Each animal was anesthetized by 3 % isoflurane, surgery was performed under 3% isoflurane and 95 % oxygen. The IPTT-tag was implanted subcutaneously into the classical brown adipose tissue (BAT) depot, using a standard IPTT-tag injector (supplied by the manufacturer). Post-mortem verification ensured the tag was in contact with the BAT. The iButton was implanted in the abdominal cavity with the use of laparotomy. At the site of laparotomy a subcutaneous injection of lidocaine mixture (lurocaine/bupivacaine, 2.5mg/kg each) was administered. Subcutaneous inject of AINS meloxicam (2mg/kg) was performed while the animal is anesthetised, and following surgery meloxicam (metacam^®^ buvable 1.5 mg / ml, dose 1 mg / kg) was added to drinking water 3 days post surgery. Drinking was monitored in the animals post-surgery establishing that they were drinking normally. The reason for administration in the drinking water is that the European hamster is an extremely aggressive therefore subcutaneous administration was not an option. The animals were extensively monitored after surgery and allowed to recover for two weeks.

### Behaviour and temperature monitoring

To keep track of individual torpor-arousal patterns, behavioural recordings were made twice per day; 1 to 2 hours after lights on and 9-10 hours after lights on. We defined torpid as a ventilation frequency (VF) <10 per minute, curled up position in hibernacula, immobile, unresponsive and an IPTT reading <13.5□. Signs of arousal were; increased IPTT tag temperature and increased ventilation frequency, at which point we began IPTT monitoring until the animal was fully awake, mobile and a euthermic body temperature was achieved, duration of monitoring varied but on average was approximately 5 hours. From the beginning of arousal IPTT tag temperatures were taken every two minutes, and ventilation frequency was counted at regular intervals for one minute. Shivering was defined as the first muscle contractions observed from the beginning of arousal monitoring. The contractions are usually observed in the scapular region, as soon as a contraction was observed we manually scanned the IPPT tag to note the BAT temperature. IButton core temperature recordings were collected post-mortem and time matched with the calibrated IPTT measurements. Graphpad Prism v8.0 was used for statistical analysis and data plotting.

### Statistical analyses

Graphpad Prism v8.0 was used for statistical analysis and data plotting. Assymetrical sigmoid curve fitting was used to analysis the temperature data from both the BAT and core body. We defined a mean baseline temperature as 0% aroused (9.8□) and euthermy as 100% aroused (37□), allowing us to establish 23.5□ as the 50% mark around which all arousal events could be synchronised and therefore plotted together to fit a best fit regression line to the linear phase of the sigmoid curve determine mean maximum re-warming rate (RWR_MAX_), and the acceleration by analysing the derivatives of the exponential and asymptotic phases of the sigmoid curve. The exponential phase and the asymptotic phase can be defined as 1 ≤ y 0 ≤RWR_MAX_, i.e. the period of time where rewarming rate is faster or equal to 1°C *h −1 but slower or equal to RWR_MAX_. This can be used as a measure of how fast RWR_MAX_ is achieved after initiation of arousal. One way ANVOVA was used to determine significant increases in ventilation rate over the arousal event. Person correlation was used to determine the correlation between ventilation and T_BAT_. Confidence intervals (CI) are given were appropriate.

## Abbrevations

BAT: brown fat
UCP1: uncoupled protein 1
ATP: Adenosine triphosphate
T_BAT_: Brown fat temperature
Tb: core body temperature
IPTT: Implantable Programmable Temperature Transponder
VF: ventilation frequency
RWR_MAX_: maximum re-warming rate

## Declarations

### Ethics approval and consent to participate

The experimental procedure was validated by a local ethical committee and further validated by the Ministry of Higher Education, Research and Innovation (APAFIS#21424-2019070219421923 v3).

### Consent for publication

Not applicable

### Availability of data and material

The datasets used and/or analysed during the current study are available from the corresponding author on reasonable request.

### Author contributions

**FM** - Experimental design, analysed the data, collected samples, prepared figures, and revised manuscript. **VJM** – Experimental design, analysed the data, collected samples, prepared manuscript. **BB** – performed surgeries, technical expertise, revised the manuscript. **DGH** - conceived and designed the experiments, supervised, and revised the manuscript. **VS** - conceived and designed the experiments, supervised, provided funding and revised the manuscript. **SHW**– conceived and designed the experiments, supervised, provided funding and prepared the manuscript. All authors read and approved the manuscript.

## Acknowledgements

The authors thank all of the animal staff at the Chronobiotron facility, especially Dominique Ciocca and Nicolas Lethenet, for their expert care of their research animals and technical assistance.

